# Arabidopsis INHIBITOR OF GROWTH 2 (ING2) promotes flowering by regulating NuA4-dependent histone H4 acetylation levels at *FT* and *SOC1* chromatin loci

**DOI:** 10.1101/2025.03.10.642062

**Authors:** Javier Barrero-Gil, Alfonso Mouriz, Raquel Piqueras, Yingnan Tian, Juan A López, Jesús Vázquez, Pedro Crevillén, José A. Jarillo, Manuel Piñeiro

## Abstract

INHIBITOR OF GROWTH (ING) proteins are chromatin readers that bind trimethylated histone H3 lysine (K) 4 (H3K4me3) marks and associate with either histone acetyltransferase or deacetylase complexes to activate or repress gene transcription, respectively. In plants, there are two types of ING proteins named ING1 and ING2. Here we report that Arabidopsis ING2 associates with multiple subunits of the histone H4 acetyltransferase complex NuA4, controls genome-wide levels of histone H4 acetylation (H4ac) and regulates different developmental processes including the initiation of flowering. Our data indicates that ING2 biological functions are largely independent of ING1 activity. We find that ING2 is recruited to the chromatin of key floral integrators such as *FT* and *SOC1*, and is required for their timely activation by modulating histone H4ac levels at these loci. Besides, our observations reveal a genetic interaction between *ING2* and genes encoding relevant proteins for H3K4me3 or H2A.Z deposition, suggesting that ING2 might represent a hub for potential crosstalk between histone H4ac and these histone modifications/variants.

## Introduction

Histones are core components of chromatin and play a fundamental role in DNA functional biology. These proteins determine chromatin compaction and accessibility (Luger et al., 1997), which profoundly affects DNA transcription and replication. Histone post-translational modifications (PTMs) have been shown to be instrumental for such processes (Strahl and Allis, 2000; Jenuwein and Allis, 2001). Histone PTMs are recognized by specific proteins, known as histone readers (Kutateladze, 2011), to trigger the recruitment of protein complexes at specific genomic loci conveying the information contained in histone PTMs (Bannister and Kouzarides, 2011).

The plant homeodomain (PHD) finger is a zinc-binding domain of 50 to 80 amino acids (Mouriz et al., 2015) that enables reader binding to di- and tri-methylated histone H3 lysine 4 residues (H3K4me2/me3), landmarks of transcriptionally active genes (Quan et al., 2023). Among the best known PHD-containing readers are the highly conserved INHIBITOR OF GROWTH (ING) proteins (He et al., 2005), which are characterized not only by their capacity to bind methylated histone H3 but also phosphoinositol monophosphates, and to regulate gene expression through the recruitment of histone acetyltransferases (HAT) and deacetylases (HDACs) complexes to activate or repress gene expression respectively (Soliman and Riabowol, 2007). The ING family comprises five members in mammals (ING1-5), which play crucial roles in DNA damage repair, cell apoptosis and tumour suppression (Taheri et al., 2022). However, phylogenetic analysis indicates that plant ING proteins group in two clades, each one representing ING1-like and ING2-like proteins (Jaudal et al., 2022). The genomes of Arabidopsis (Lee et al., 2009) and Medicago (Jaudal et al., 2022) contain a representative of each group and both display the capacity to bind the active transcription epigenetic mark H3K4me3 *in vitro* (Lee et al., 2009; Zhao et al., 2018). Plant ING proteins may associate with several chromatin modifying complexes. In Arabidopsis, ING1 was recently shown to recruit a plant-specific GCN5 (GENERAL CONTROL NON-DEPRESSIBLE 5)-containing complex called PAGA that catalyzes the acetylation of histone H3 (Wu et al., 2023). Also in Arabidopsis, proteomic studies and protein-protein binding assays have shown that ING2 proteins interact with different chromatin remodeling factors including histone deacetylases like HDC1 (Perrella et al., 2016) and NuA4-C (for nucleosome acetyltransferase of H4) subunits (Tan et al., 2018; Barrero-Gil et al., 2022; Bieluszewski et al., 2022). However, whether plant ING2 proteins are core subunits of these complexes remains unclear, and no proteomic studies using ING2 as a bait have been reported so far. ING2 promotes flowering in Medicago (Jaudal et al., 2022), though the exact mechanism remains to be elucidated. Interestingly, NuA4-C is known to promote flowering in Arabidopsis through H2A.Z (Crevillen et al., 2019) and H4 acetylation (Xiao et al., 2013; Barrero-Gil et al., 2021). Therefore, it is tempting to speculate that ING2 may regulate flowering through the modulation of NuA4-C-histone acetylation activity. In addition, whether ING2 recruits additional protein complexes remains an open question.

In this work, using Arabidopsis as genetic model, we take unbiased proteomics approaches to unveil ING2-bound proteins, what is the relationship between the two Arabidopsis ING proteins, what is the physiological role of ING2 and which is the molecular mechanism that enables ING2 to regulate flowering. The results demonstrate that ING2 is part of NuA4-C, and acts modulating genome-wide histone H4ac levels and controlling different developmental responses including flowering time. Our results argue for distinct epigenetic mechanisms mediating the function of the two plant types of ING readers in regulating gene expression. In particular, Arabidopsis ING2 controls flowering time through the regulation of H4ac levels at chromatin loci encoding key flowering time regulators.

## Results

### ING2 is co-purified along with most subunits of NuA4-C

We followed AP-MS approaches to identify the proteins that can co-purify with the ING2 protein in aerial tissues from 12 day-old seedlings grown under long days (LD) and 22°C. We identified among others, peptides corresponding to 7 out of the 12 subunits that constitute the plant NuA4-C (Espinosa-Cores et al., 2020), including proteins from the Piccolo, YEATS and core modules (Figure 1A, Supplemental Figure 1, Supplemental Table 1), while no peptide corresponding to any subunit from the TINTIN module copurified with ING2. These results together with our previous data (Barrero-Gil et al., 2022) and data published by other groups (Bieluszewski et al., 2015; Tan et al., 2018; Bieluszewski et al., 2022; Zhou et al., 2022; Zheng et al., 2023) firmly establish ING2 as a bona-fide member of the NuA4-C (Supplemental Fig.1). Remarkably, we failed to detect peptides from any HDAC complex subunits (Supplemental Table 1). We used the protein-protein interaction database STRING (https://string-db.org/) to further analyze ING2 co-purifying proteins, finding protein networks related to pre-mRNA processing factor 19 (PRP19) (Chanarat and Strasser, 2013), Mediator (Buendia-Monreal and Gillmor, 2016) and the NuA4 complexes (Supplemental Table 1, Figure 1B). A functional enrichment analysis of this network revealed terms related to H4 acetylation and mRNA splicing.

**Figure 1:**
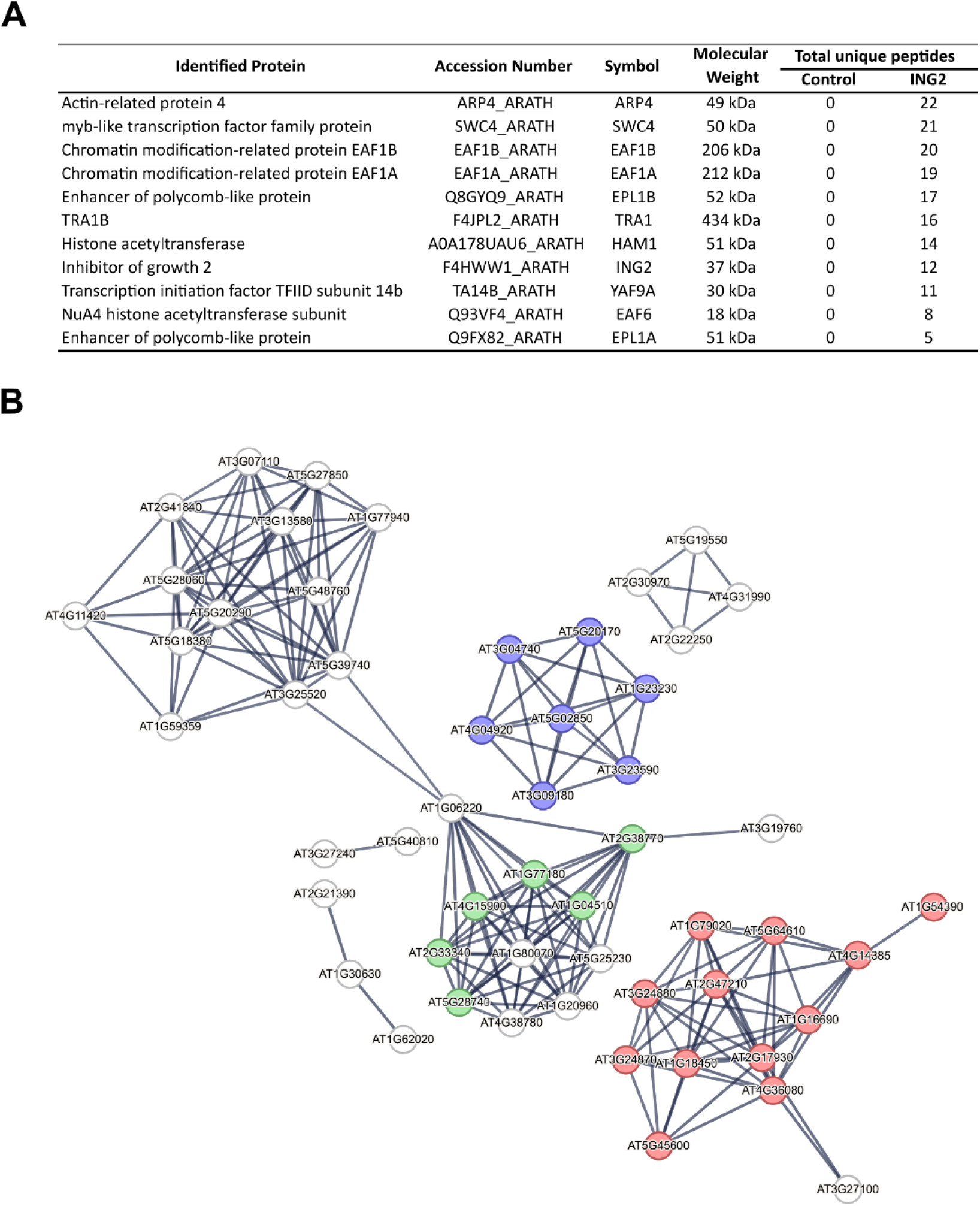
ING2 interacts with NuA4 complex subunits. (A) List of NuA4-C subunits identified among proteins co-purified with the ING2 protein in this work. (B) Functional protein networks identified among ING2-copurifying proteins. PRP19, Mediator and NuA4 complexes have been highlighted in green, purple and red respectively.

### ING2 regulates flowering time and additional developmental processes

To unveil the functions carried out by the ING2 protein, we obtained from the Arabidopsis NASC germplasm collection two T-DNA mutant lines, GABI_166D07 (*ing2-1*) and GABI_909H04 (*ing2-2*), that were confirmed to be knockout (KO) mutants for *ING2* (Supplemental Figure 2A-B). Loss-of-function alleles of *ING2* resulted in a delayed flowering time under LD (Figure 2A), in line with observations made in Medicago (Jaudal et al., 2022). Intriguingly, *ing2* mutant plants grown under short days (SD) did not flower, entering senescence before bolting (Figure 2B). Additionally, over-expression of *ING2* complemented the late flowering phenotype observed in the *ing2* KO mutants (Supplemental Fig 2), confirming that ING2 is a positive regulator of the floral transition. Further analysis showed that the loss of *ING2* function also affects chlorophyll accumulation (Figure 2C) and flower architecture, altering the number of petals (Figure 2D).

**Figure 2:**
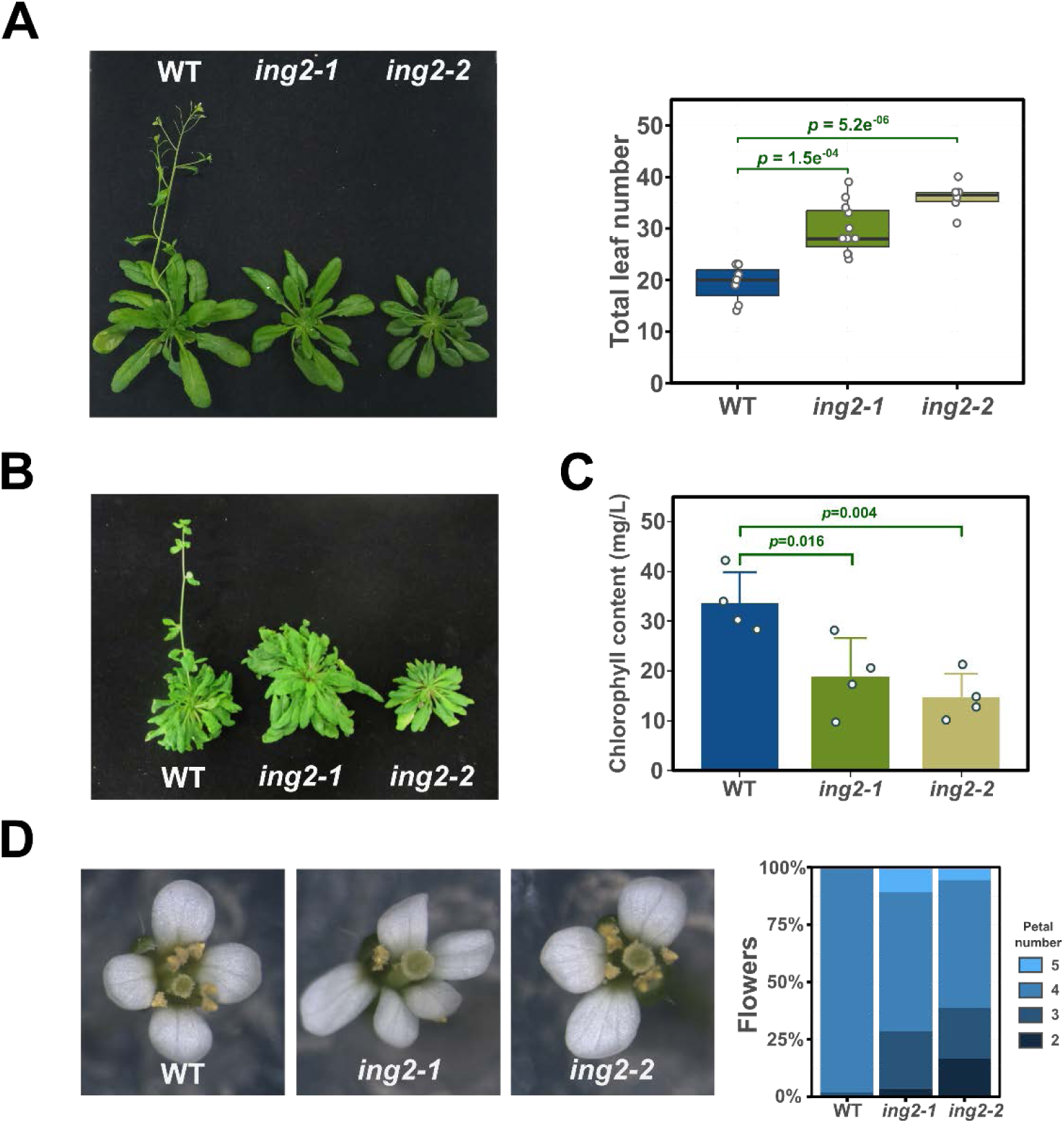
Arabidopsis *ING2* is involved in the promotion of flowering and additional developmental processes. (A-B) Flowering time in *ing2* mutant plants under long day (A) or short day (B) photoperiod. Quantification under SD is not feasible since most of the plants do not flower in these conditions. (C-D) Analysis of chlorophyll content (C) and flower development (D) in *ing2* mutant plants. Statistical significance is indicated providing *p*-value in a Dunnet’s test (A and C).

### ING2 biological functions are largely independent of ING1 activity

The Arabidopsis genome includes two closely related ING proteins (Lee et al, 2009). To study the functional relationship between these two genes, we isolated *ing1* loss of function mutants (Wu et al., 2023) to generate *ing1 ing2* double mutant plants and compared flowering time with *ing1* and *ing2* single mutant plants. The results reveal opposite effects of the loss of *ING1* and *ING2* function on the determination of flowering time as *ing1* mutant plants tend to flower slightly earlier than wild-type (WT) under LD, while *ing2* mutants flower much later (Figure 3A). Double *ing1 ing2* mutant plants also display a late flowering phenotype similar to the *ing2* single mutant plants. Under SD, the *ing1* mutant plants did not display obvious alterations in flowering time while both *ing2* and *ing1 ing2* double mutant plants failed to flower (Figure 3B). In addition, we observed that *ing2* mutants showed smaller rosettes in these conditions regardless of the presence of an active ING1 protein (Figure 3C-D). Moreover, *ing1* mutants did not show altered levels of chlorophyll in clear contrast to *ing2* and *ing1 ing2* mutants that showed a similar reduction in chlorophyll content under LD photoperiod (Figure 3E). Finally, we observed that the *ing1 ing2* double mutant produced shorter siliques under LD photoperiod (Supplemental Figure 4). We measured silique length in single and double *ing* mutants, unveiling a synergistic effect of these mutations on the length of siliques (Supplemental Figure 4). These results indicate that ING2 function in the regulation of flowering time, rosette size or the chlorophyll content is not significantly affected by the loss of *ING1* function, while a more complex relationship between *ING* genes can be observed in the regulation of other developmental traits like fruit development, where loss-of-function of both genes results in an enhancement of the phenotypic alterations observed.

**Figure 3:**
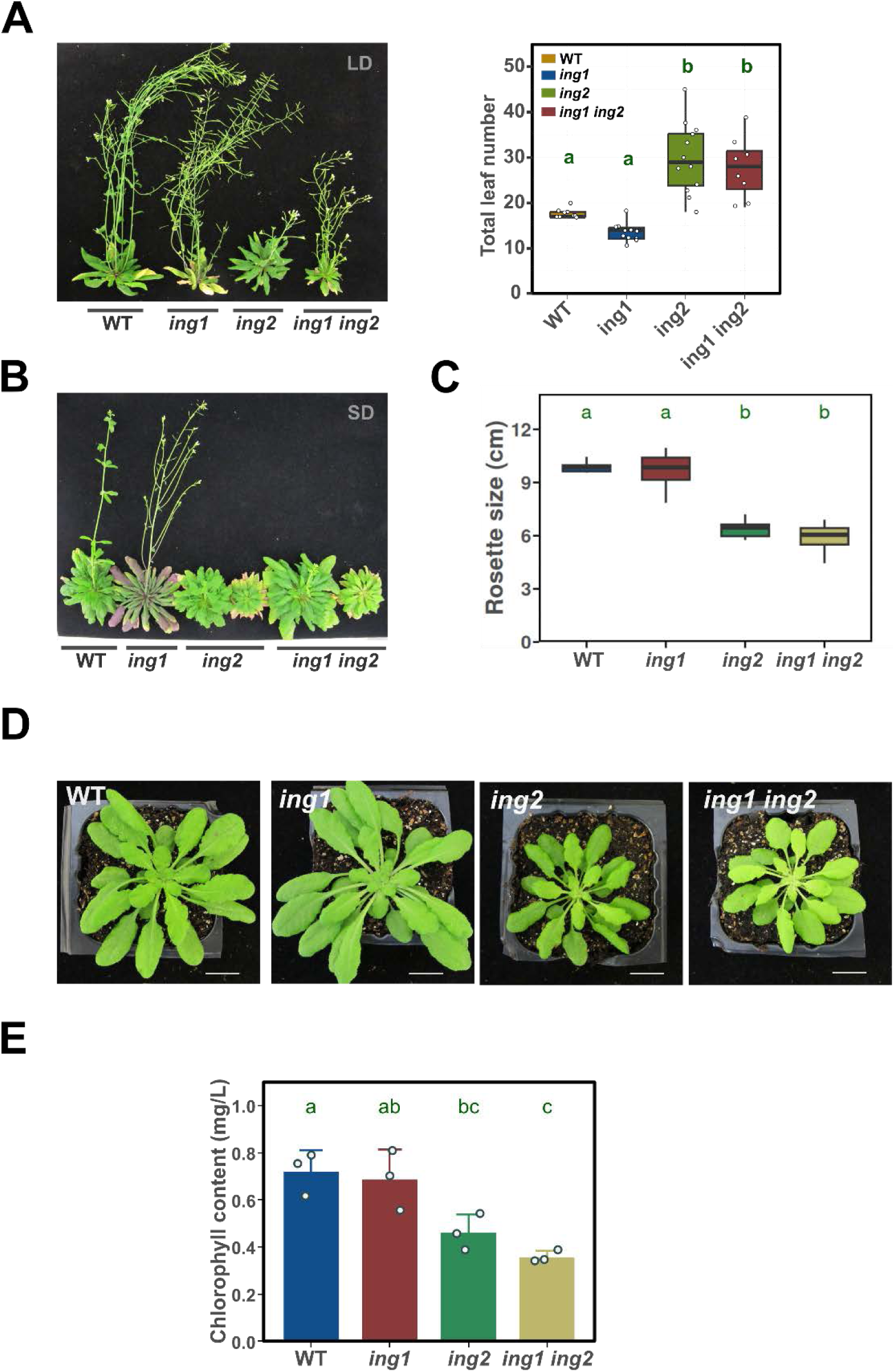
Developmental alterations in *ing2* mutant plants are independent of *ING1* function. (A-B) Genetic relationship of *ING1* and *ING2* genes in the determination of flowering time under long- (A) or short-day (B) photoperiod. (C) Quantification of rosette diameter in plants grown under SD. (D) Representative plants of the indicated genotypes grown under non-inductive photoperiodic conditions. (E) Determination of chlorophyll content in 10 day-old seedlings grown under LD. Statistical significance in one-way ANOVA tests is indicated by different letters.

To further study the relationship between ING1 and ING2 proteins we compared global gene expression profiles in WT, *ing* single and double mutant seedlings. We detected 195, 354 and 956 upregulated genes, and 40, 90 and 310 downregulated genes in the *ing1*, *ing2*, and *ing1 ing2* double mutant, respectively (Figure 4, Supplemental Table 2). Interestingly, the hierarchical clustering analysis grouped *ing2* and *ing1 ing2* together (Figure 4A). Furthermore, in a principal component analysis (PCA) the transcriptomic profile of *ing1* mutant plants is not clearly separated from that of WT plants in a graphical representation using the first two principal components (Figure 4B). In contrast, the transcriptomic profiles of *ing2* single and double mutant plants are clearly separated from that of WT and *ing1* plants (Figure 4B). Venn diagrams showed a significant but limited overlap of genes regulated by ING1 and ING2, particularly among downregulated genes (Figure 4C). A comparison of differentially expressed genes (DEG) in *ing* mutants with ENHANCER OF POLYCOMB-LIKE 1 (EPL1) (Barrero-Gil et al., 2022) and GCN5-regulated genes (Wu et al., 2023), representing NuA4-C or PAGA-dependent genes respectively, reveals that misregulated genes in *ing2* mutants are more enriched in NuA4-C-dependent genes than in PAGA-dependent genes, while the reverse is true for *ing1* mutants (Figure 4D). A gene ontology (GO) enrichment analysis reveals different specific and shared biological terms among DEGs in *ing* mutants (Supplemental Figure 5, Supplemental Table 2). All in all, genetic and transcriptomic analysis suggest that Arabidopsis ING proteins co-regulate a very limited set of genes, consistent with both proteins being part of different chromatin remodeling complexes.

**Figure 4:**
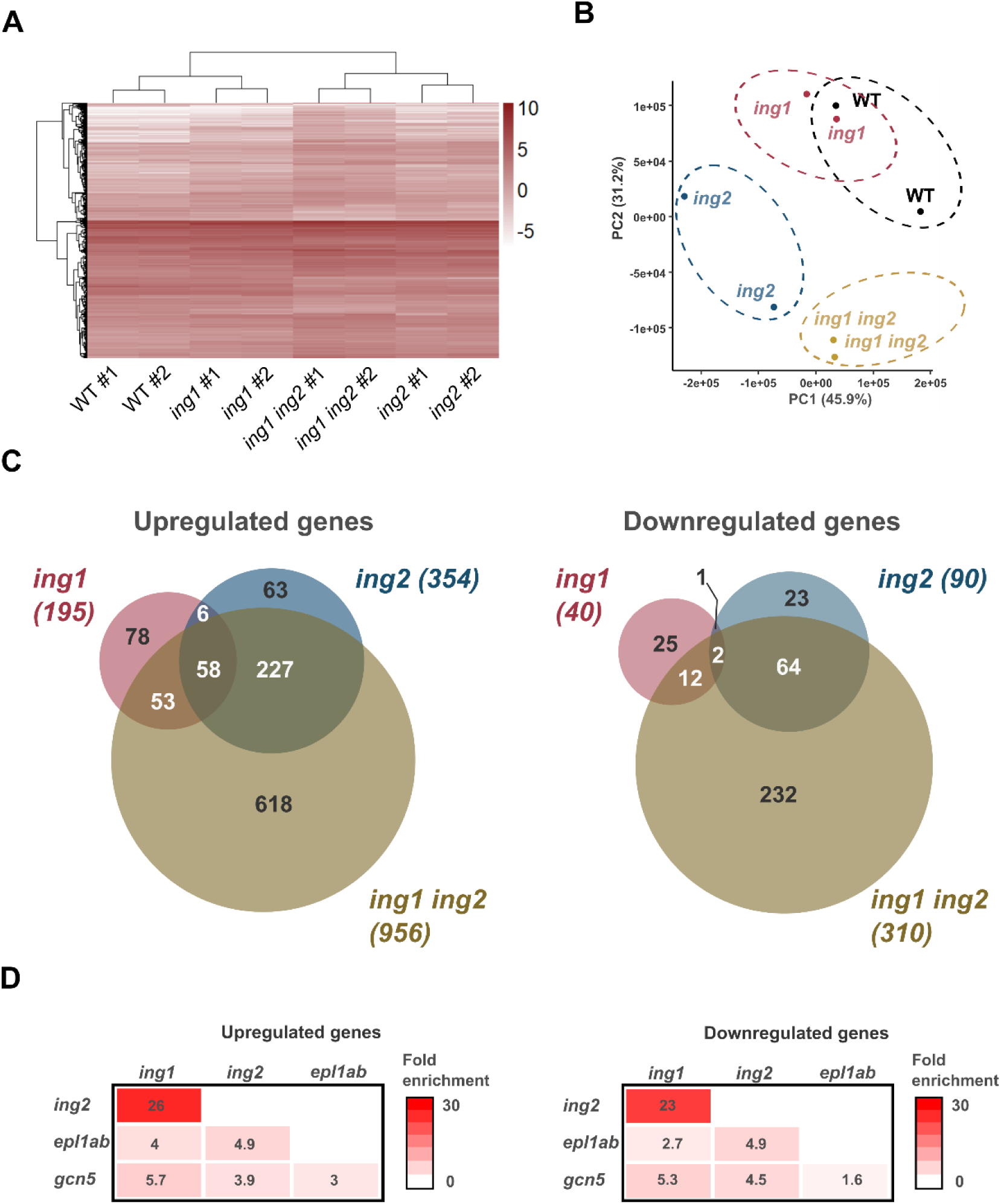
ING1 and ING2 mostly regulate different gene sets. (A) Heatmap showing expression level of DEGs in *ing* mutants. (B) Principal component analysis of expressed genes in the indicated genotypes. (C) Overlap among DEGs in *ing* mutants. (D) Heatmap showing the degree of overlap between ING-regulated genes and genes regulated by NuA4-C (*epl1ab*) (Barrero-Gil et al., 2022), and PAGA-C (*gcn5*) (Wu et al., 2023). Fold enrichment for each pairwise comparison is indicated by color scale and the number on each panel.

### ING2-mediated regulation of the floral transition requires di- and tri-methylated H3K4 marks

Previous work has shown that Arabidopsis ING2 can bind the H3K4me2/me3 activating marks (Lee et al., 2009). Trithorax protein complexes are responsible for methylating histone H3K4 residues in eukaryotes (Avramova, 2009). The Arabidopsis genome includes ten Trithorax proteins (Tamada et al., 2009), and ARABIDOPSIS TRITHORAX1 (ATX1)/ SET DOMAIN GROUP 27 (SDG27) (Pien et al., 2008) and ARABIDOPSIS TRITHORAX-RELATED7 (ATXR7)/SDG25 (Tamada et al., 2009) have been shown to regulate flowering time. We decided to explore genetic relationships of *ING2* with the genes encoding these two proteins by crossing the corresponding single loss-of-function mutant plants and analyzing the floral transition in the double mutant. The results show that the loss of function of either *ATX1* or *ATXR7* reverted the late flowering phenotype displayed by *ing2* mutant plants (Figures 5A-B), indicating that *ING2* function is dependent on histone H3K4me2/me3 levels.

**Figure 5:**
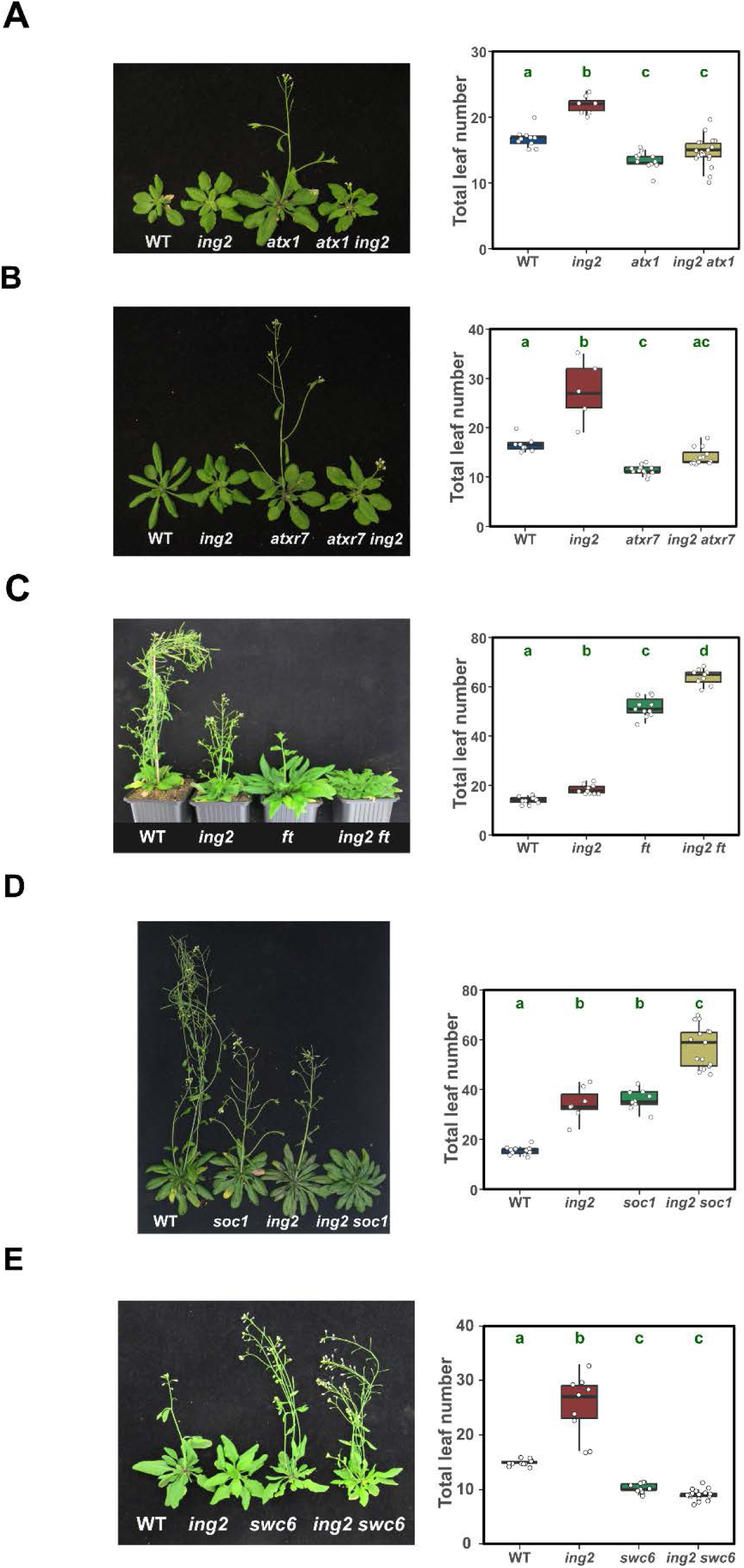
Genetic interactions provide insights into ING2 function in the regulation of flowering time. Flowering time was determined in single and double mutants deficient in the activities of *ING2* and *ATX1* (A), *ATXR7* (B), *FT* (C), *SOC1* (D) or *SWC6* (E) genes. Left panels show representative plants of the indicated genotypes. Right panels show flowering time scored as the number of total leaves. Plants were grown under LD photoperiodic conditions.

### The role of ING2 in flowering time control does not rely on the function of a single floral integrator and requires SWR1-C activity

Our data show that the ING2 protein interacts with a number of NuA4-C subunits and that it is necessary to promote flowering (Figures 1 and 2). In previous works, NuA4-C was shown to regulate the initiation of flowering through the acetylation of histone H4 at *FT* and *SOC1* chromatin (Xu et al., 2014; Barrero-Gil et al., 2021) and H2A.Z at *FLC* chromatin (Crevillen et al., 2019). To address the mechanism by which ING2 regulates flowering, we crossed the *ing2* mutant with *flowering locus t* (*ft*) (Yoo et al., 2005) and *suppressor of overexpression of constans1* (*soc1*) (Lee et al., 2000) mutants, affected in the central floral integrators *FT* and *SOC1*, respectively, and quantified their flowering time under a LD photoperiod regime (Figures 5C-D). The results show an additive effect of *ing2* and *ft* and *soc1* mutations, indicating that either ING2 role in flowering does not depend on a single floral integrator or such role is completely independent on *FT* and *SOC1* function.

Recent evidence suggests that NuA4-C may regulate the levels of the histone variant H2A.Z at certain genomic locations (Bieluszewski et al., 2022). To investigate a possible relationship between ING2 function and the deposition of the histone variant H2A.Z, we crossed the *ing2* mutant with the *swr1 complex subunit 6* (*swc6*) mutant that is defective on the Swi2/Snf2-related chromatin remodeling complex SWR1 (SWR1-C) function and hence on H2A.Z deposition on chromatin (Lazaro et al., 2008). The *swc6* mutation causes plants to flower earlier than WT plants. The results show a clear epistatic relationship between *SWC6* and *ING2*, indicating that *ING2* function is dependent on SWR1-C activity (Figure 5E). Altogether, our genetic analyses indicate that ING2 proteins integrate information relayed by histone H3K4me3 epigenetic marks, controlling flowering time possibly through several floral integrators. Such control is largely dependent on the activity of SWR1-C.

### ING2 promotes genome-wide H4 acetylation but not H2A.Z chromatin deposition

Since the ING2 protein interacts with many NuA4-C subunits (Figure 1), we anticipated that ING2 might contribute to the activity of this complex. To tackle this issue, we sequenced libraries prepared from chromatin immunoprecipitated (ChIP-seq) with antibodies raised against poly-acetylated histone H4 or unmodified histone H4 as a control. We also performed a ChIP-seq experiment to analyze chromatin H2A.Z levels, since previous observations proposed that the NuA4-C promotes the deposition of this histone variant (Bieluszewski et al., 2022) and our genetic evidence showed an epistatic relationship between *ING2* and genes encoding proteins involved in H2A.Z deposition (Figure 5). The results show a global decrease in histone H4 acetylation levels in *ing2* chromatin compared to WT but no significant changes in chromatin H2A.Z levels (Figure 6A). Intriguingly, we also observed a reduction of H4 occupancy upstream and downstream of the transcriptional start site (TSS) (Supplemental Figure 6A). To check if this circumstance could challenge our interpretation of the data, we normalized the H4ac and H2A.Z signals by H4 occupancy level reaching the same conclusions, i.e., ING2 promotes H4 acetylation but does not influence H2A.Z occupancy (Supplemental Figure 6B). Next, we asked whether the altered levels of histone H4ac in the *ing2* mutant are associated with changes in gene expression. To do that, we averaged H4ac signal over a region spanning 500 base pairs downstream of the TSS for each gene and calculated the change in H4ac as the log2 difference between the *ing2* mutant and WT. We sorted genes according to H4ac change, divided them in percentiles and for each percentile, we averaged the H4ac change and gene expression level, which was derived from our transcriptomic analysis of *ing* mutants. Spearman rank correlation analysis revealed a significant association between H4ac and gene expression change in *ing2* mutant compared to WT (Figure 6B). Next, we performed a differential peak enrichment analysis on our ChIP-seq data. The results show that the overwhelming majority of differentially acetylated genes correspond to genes with lower H4ac in the *ing2* mutant (Figure 6C). A gene ontology analysis on the genes hypoacetylated in the *ing2* mutant revealed an enrichment of terms related to photosynthetic functions including plastid translation, chlorophyll biosynthesis and chloroplast fission (Figure 6D). Interestingly, among the genes identified with lower H4ac levels we found diverse flowering integrators such as *SOC1* (Samach et al., 2000), *GIGANTEA* (Fowler et al., 1999) and *CONSTANS* (Putterill et al., 1995) but could not identify *FT* (Kardailsky et al., 1999), probably because of the low H4ac signal detected within this genomic region (Figure 6E, Supplemental Table 3). We sought to confirm the results for *FT* and *SOC1* genes by performing ChIP-PCR. Remarkably, besides obtaining confirmatory data on the low H4ac status of *SOC1* chromatin, we also found a moderate but significant reduction of *FT* locus H4ac level in *ing2* mutants (Figure 6F).

**Figure 6.**
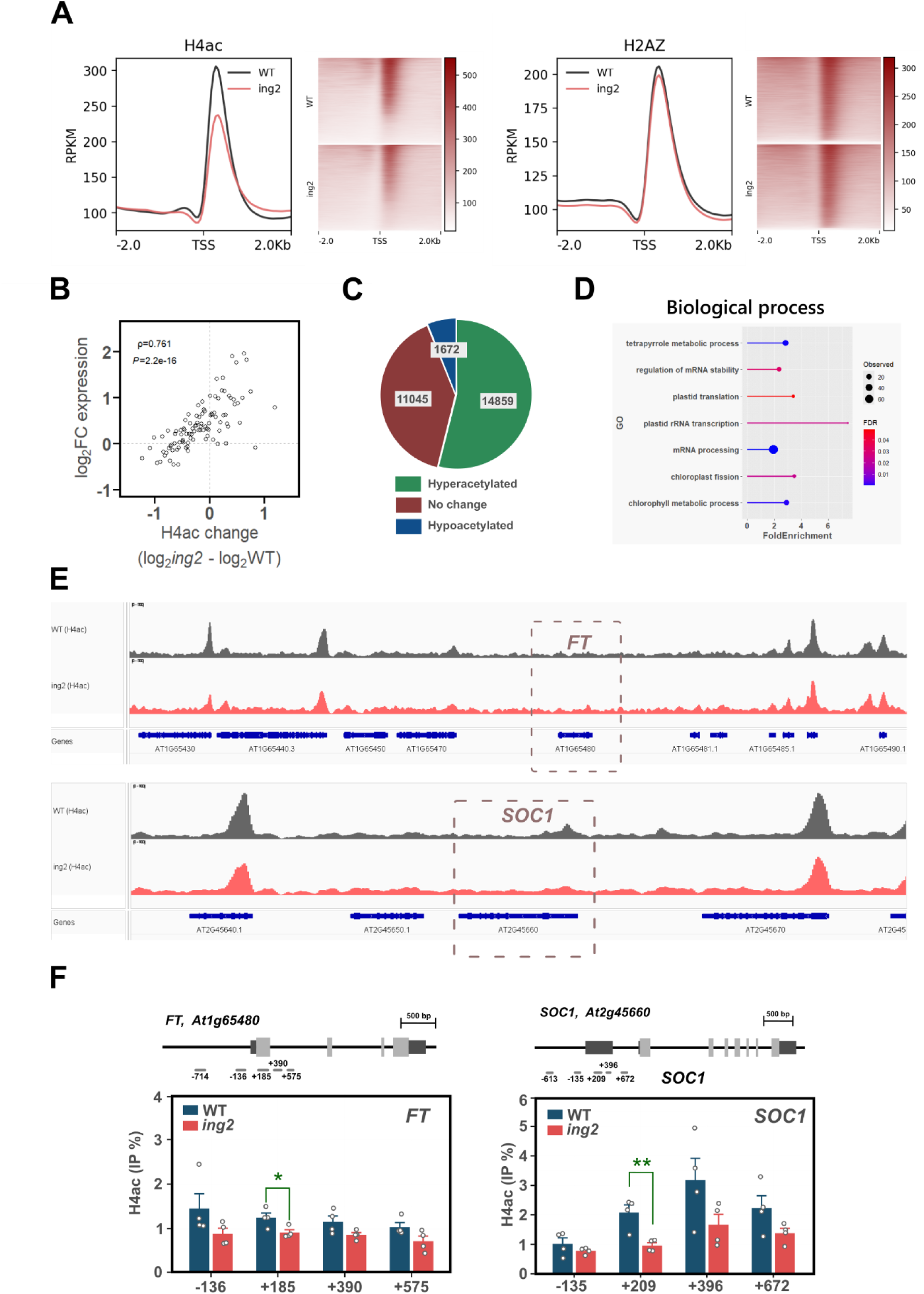
ING2 promotes global H4 acetylation but does not influence H2A.Z deposition. (A) Metaplot and read density maps showing H4ac and H2A.Z signal level and distribution pattern in WT and *ing2* mutant plants. (B) Spearman rank correlation analysis between changes in H4ac levels and gene expression observed in *ing2* mutant plants. Spearman coefficient and *p*-value are indicated in the inset within the scatter plot. (C) Number of hypoacetylated (green) and hyperacetylated (blue) peaks in *ing2* chromatin. (D) Enrichment analysis of gene ontology terms associated with the hypoacetylated peaks identified in (C). (E) H4ac profiles of wild type (grey) and *ing2* (red) in genomic regions centered on *FT* and *SOC1* genes. (F) H4ac levels at *FT* and *SOC1* genomic loci as determined by ChIP-qPCR experiments. Tested amplicons are indicated over each graph and significant differences in two-sided t-tests are indicated with asterisks.

### ING2 regulates flowering time activating *FT* and *SOC1* transcription

To clarify if the low H4ac status on *FT* and *SOC1* chromatin in *ing2* mutants is correlated with a lower expression of these genes, we performed a time-course gene expression analysis of these floral integrator genes in the *ing2* mutant (Figure 7A). The results obtained show a clear decrease in the activation of both flowering activators. To confirm that these genes might be direct targets of *ING2* regulation, we used in ChIP experiments the transgenic line that overexpresses a myc-tagged construct of ING2 complementing the flowering delay observed in *ing2-2* mutant plants (Supplemental Figure 3), previously employed for our AP-MS analysis (Figure 1). Quantitative PCR amplification showed an enrichment in the myc-ING2 immunoprecipitated fraction of selected amplicons in *FT* and *SOC1* loci coinciding with reported histone H3K4me3-enriched regions (Zhu et al., 2024). These observations indicate a direct binding of ING2 to these master regulator genes of flowering (Figure 7B). In summary, our results indicate that ING2 directly regulates the transcript levels of *FT* and *SOC1* floral regulators by controlling their histone H4 acetylation levels.

**Figure 7:**
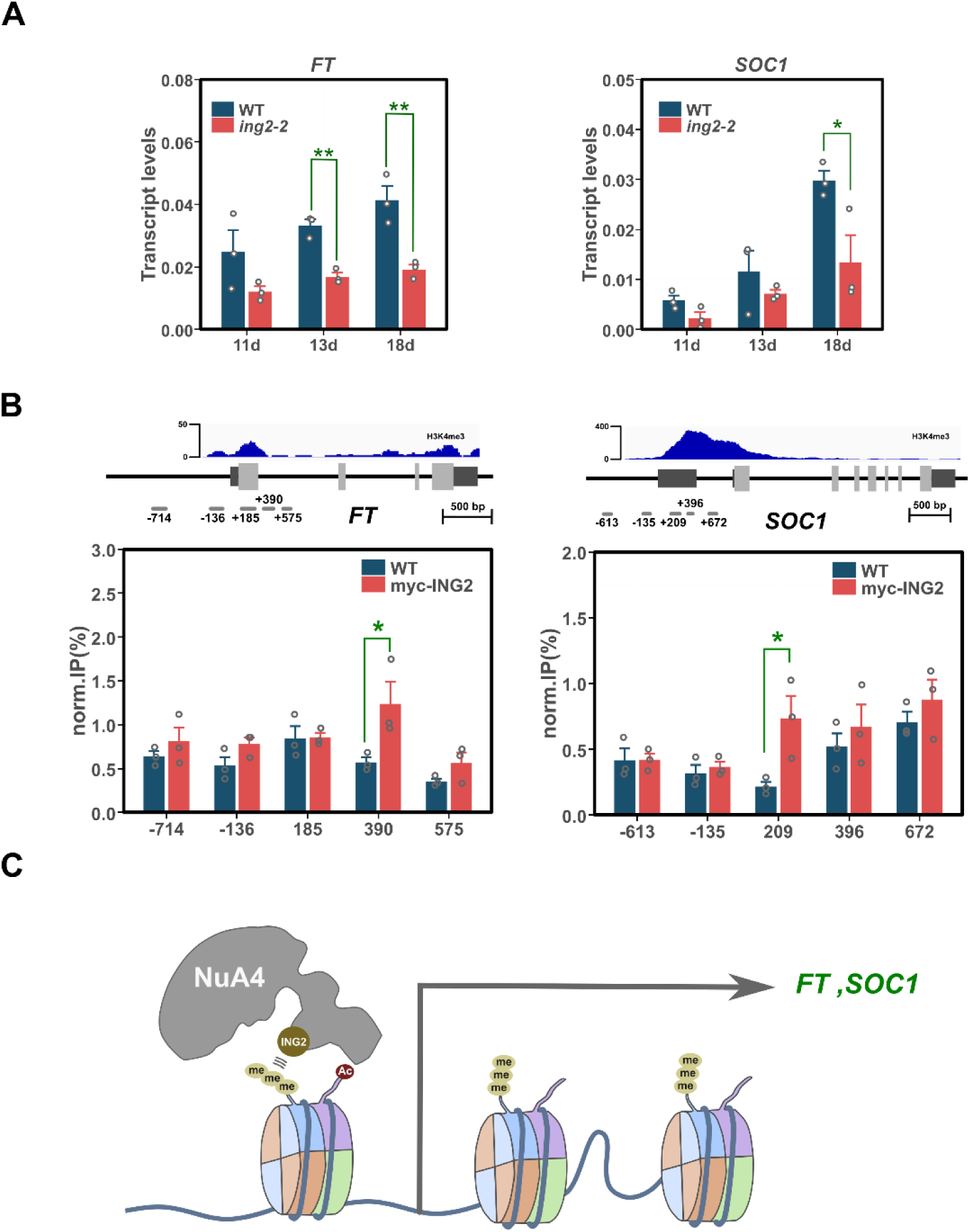
ING2 protein binds *FT* and *SOC1* chromatin to regulate H4ac level promoting the activation of the floral integrator genes. (A) Developmental expression analysis of floral integrators in WT and *ing2* mutant. (B) ChIP-qPCR experiments in WT and transgenic *35S-myc-ING2* plants. The data show the immunoprecipitated fraction recovered from *FT* or *SOC1* chromatin after immunoprecipitation with the anti-myc antibody. Coverage data for H3K4me3 in WT is indicated as a reference above gene structure. (C) Hypothetical working model on ING2 function. As part of NuA4-C, ING2 subunit likely enables H3K4me3 reading and participates in the modulation of H4ac levels in floral integrator target genes *FT* and *SOC1*. A similar molecular mechanism might mediate the involvement of ING2 in the regulation of additional developmental processes that are altered in the *ing2* mutant by modulating acetylation levels in the chromatin of other target genes. Bars indicate the average of three biological replicates and error bars denote SEM. Asterisks denote *p*<0.05 in two-sided t-tests

## Discussion

To dissect the function of plant PHD-containing ING proteins, we have focused on the Arabidopsis ING2 protein, identifying co-immunoprecipitated proteins, analyzing the relationship with Arabidopsis ING1, and characterizing the physiological and molecular function of ING2. Our results establish ING2 as a bona-fide component of NuA4-C in plants, regulating a variety of developmental processes, including flowering time through the modulation of histone H4 acetylation at loci encoding key floral integrators such as *FT* and *SOC1*.

Our AP-MS analysis detected peptides corresponding to 7 out of 12 subunits that putatively constitute the plant NuA4-C (Espinosa-Cores et al., 2020). Together with data from additional AP-MS studies using different NuA4-C subunits as baits (Bieluszewski et al., 2015; Tan et al., 2018; Bieluszewski et al., 2022; Zhou et al., 2022; Zheng et al., 2023) as well as our previous protein-protein interaction results (Barrero-Gil et al., 2022) these observations provide solid evidence for ING2 as a core member of the plant NuA4-C (Figure 1A; Supplemental Figure 1). Unexpectedly, this experiment did not identify peptides corresponding to proteins such as Histone Deacetylase Complex 1 (HDC1), Sin3A-Associated Protein 18 (SAP18) or SHORT LIFE (SHL) that showed capacity to bind ING2 in tobacco transient co-expression experiments (Perrella et al., 2016). Similarly, the Medicago ING2 protein does not appear to interact with them either in yeast-two-hybrid assays (Jaudal et al., 2022). It is possible that the experimental conditions used in our study favor ING2 interaction with the NuA4-C. Alternatively, perhaps the recombinant *ING2* gene construct we used, though able to complement the phenotypic defects in *ing2* mutant plants, affects the capacity of tagged ING2 to effectively interact with HDACs. Intriguingly, we also identified in our AP-MS analysis peptides corresponding to proteins involved in pre-mRNA-splicing (Chanarat and Strasser, 2013) as well as proteins from the Mediator complex (Buendia-Monreal and Gillmor, 2016), (Figure 1B), suggesting the ING2 function might be also linked to transcription initiation or mRNA splicing processes, an issue that will require further investigation.

Similar proteomic approaches using ING1 as bait have failed to identify NuA4-C subunits (Wu et al., 2023). Conversely, AP-MS studies using different NuA4-C subunits as baits have repeatedly failed to identify peptides corresponding to the ING1 protein. These observations suggest that ING1 and ING2 proteins might enable H3K4me3 reading in different chromatin modifying complexes. In this context, while the Arabidopsis *ing2* mutant is characteristically late flowering, a recent study could not identify a similar phenotype for Arabidopsis *ing1* mutant (Wu et al., 2023). We performed a genetic interaction analysis to address the relationship between both genes and found that *ING2* functions are largely independent of ING1 activity in the control of flowering time and other developmental processes. Moreover, the transcriptome regulated by ING1 is closer to the PAGA-dependent transcriptome while the ING2-dependent transcriptome is more related to NuA4-C-regulated genes. Interestingly, a phylogenetic analysis of plant genes encoding ING proteins reveals that ING1 and ING2 plant clades appeared early in the green lineage (Jaudal et al., 2022). All these observations support the notion that despite their relative sequence similarity and domain composition, ING1 and ING2 proteins likely belong to separate chromatin modifying complexes in Arabidopsis. As shown in Figure 4C, we observe a larger number of misregulated genes in the *ing1 ing2* double mutant than in the single *ing1* or *ing2* mutants. This enhanced transcriptional misregulation detected in the double mutant plants might be due to the magnifying effect that simultaneous loss of NuA4 and SAGA-like HAT complexes could have on plant gene expression. In fact, several studies in yeast have already highlighted the interplay between NuA4 and SAGA complexes in gene transcription through chromatin acetylation (Durant and Pugh, 2006; Ginsburg et al., 2009; Bruzzone et al., 2018). Further studies will be needed in plants to fully understand possible relationships between histone H3 and H4 acetylation in the regulation of gene expression.

Our physiological analyses of ING2 function revealed that this protein is involved in various developmental processes including flowering time, chlorophyll accumulation, and flower and fruit development. These phenotypic alterations are reminiscent of a loss-of-function *ING2* Medicago mutant characterized (Jaudal et al., 2022), indicating that ING2 physiological roles may be conserved in plants. Several mutant plants lacking the function of diverse NuA4-C subunits exhibit either an acceleration (Xiao et al., 2013; Bieluszewski et al., 2015; Gomez-Zambrano et al., 2018; Crevillen et al., 2019) or a delay of flowering time (Bu et al., 2014; Xu et al., 2014; Barrero-Gil et al., 2021). In our current study we observed that the ING2 protein is a promoter of flowering in photoperiod-inductive conditions and becomes absolutely essential for flowering under non-inductive conditions. We revealed that the ING2 protein activates the expression of *FT* and *SOC1* promoting histone H4ac levels at these chromatin loci. Moreover, our profile of H4ac in WT and *ing2* mutant plants demonstrates that such alterations in H4ac are not restricted to these genes but rather can be found genome-wide. Hence, we conclude that ING2 chromatin reader modulates H4ac levels and that alterations in its activity correlate with changes in gene expression.

ING proteins have been shown to specifically recognize and bind H3K4me3 *in vitro* through their PHD domain (Lee et al., 2009). To gather further insights into the activity of ING2 as a H3K4me3 reader, we analyzed whether ING2-mediated flowering time was dependent on the function of H3K4me3 writers ATX1 and ATXR7 (Pien et al., 2008; Tamada et al., 2009). The results reveal an epistatic relationship between these genes regarding the regulation of flowering time, which is consistent with a functional link between H3K4me3 deposition and ING2 activity that mediate NuA4-C acetylation on target genes bearing particular chromatin conformations or histone marks. Previous observations with the ING4 homolog, a subunit of the versatile HAT HBO1, support a crosstalk between histone H3K4 trimethylation and H3ac to attenuate cellular neoplastic transformation in metazoans (Hung et al., 2009). Finally, we have also explored a functional relationship between ING2 and SWR1-C activity (Figures 5 and 6). Our genetic interaction analysis revealed the dependence of *ing2* late flowering phenotype on SWR1-C subunits (Figure 5). Intriguingly, though previous works revealed a correlation between the NuA4-C activity mediated by EPL1 and H2A.Z global chromatin levels (Bieluszewski et al., 2022), we did not find any significant alterations in genome-wide deposition of the H2A.Z histone variant in the *ing2* mutant (Figure 6). Perhaps, ING2 might be dispensable for NuA4-C-mediated regulation of H2A.Z deposition or the affectation of global H2A.Z may require higher levels of inhibition of NuA4-C activity.

Our data support a working model (Figure 7C) in which the ING2 subunit likely enables NuA4-C H3K4me3 reading and participates in the modulation of H4ac levels on underlying genes. This activity is required for several developmental processes, mainly the initiation of flowering, as well as flower and fruit development or chlorophyll accumulation. Importantly, ING2 function is largely independent of ING1 protein and does not seem to contribute significantly to the deposition of the histone variant H2A.Z. Further study is needed to dissect the molecular basis of the genetic interaction between *ING2* and genes encoding key subunits for the SWR1-C identified in our study.

## Methods

### Plant materials, phenotypic analysis and growth conditions

All Arabidopsis lines used in this work were in Columbia-0 background. The mutant alleles *ing2-1* (GABI_166D07) and *ing2-2* (GABI_909H04) were obtained from the Nottingham Arabidopsis Stock Centre (NASC, UK). Other mutants used in this work have been previously described elsewhere: *ft-10* (Yoo et al., 2005), *soc1-2* (Lee et al., 2000), *atx1-2* (Pien et al., 2008), *atxr7*-2 (Tamada et al., 2009), *ing1-1* (Wu et al., 2023) and *swc6-1* (Lazaro et al., 2008). To generate *ing2* complemented lines we amplified *ING2* open reading frame with the primers we indicated in Supplemental Table 4 and cloned it in a pGWB18 expression vector (Nakagawa et al., 2007) that was transformed into *ing2-2* mutant plants by the floral dip method (Clough and Bent, 1998). Unless otherwise indicated, plants were grown at 22°C under LD photoperiods (16 hours of cool-white fluorescent light) with photon flux of 100 μmol m^−2^ s^−1^, in pots containing a mixture of organic substrate and vermiculite (3:1, v/v), or in Petri dishes containing 1/2x Murashige and Skoog (MS) medium supplemented with 1% sucrose where indicated, and solidified with 0.8% (w/v) plant agar.

Flowering time was determined by counting the total leaf number at the time of first flower opening (Lazaro et al., 2008) under both LD and SD photoperiods. Chlorophyll extraction was performed by overnight incubation of grounded tissue in N,N, dimethylformamide. Absorbance measurements at a wavelength of 647 and 664 nm of cleared extracts were taken on a C7100 spectrophotometer (PEAK Instruments) and chlorophyll concentration was calculated as described (Porra, 2002).

### AP-MS analysis

Arabidopsis *ing2* mutant plants complemented with the 4xmyc-ING2 construct were generated (Supplemental Figure 3). For that, the ING2 coding sequence was mobilized to the destination vector pGWB18 (Nakagawa et al., 2007), which bears the constitutive CaMV 35S promoter and a 4xMyc epitope in N-terminal, by a recombination reaction with LR clonase I (Invitrogen). The resulting clone was checked with restriction enzymes and transformed into competent *Agrobacterium tumefaciens* AGL0 cells to produce transgenic 4xmyc-ING2 *ing2* plants that fully rescued the WT flowering time and were used for AP-MS analysis. Affinity purification was performed as described (Gomez-Zambrano et al., 2018) with slight modifications. Protein extraction was performed in extraction buffer (20 mM TrisHCl pH 8, 150 mM NaCl, 2.5 mM EDTA, 33 mM β-mercaptoethanol, 10% Glycerol and 0.5% Triton X-100 supplemented with cOmplete Protease Inhibitor Cocktail (Roche, Switzerland)) from 5 g of 12 day-old seedlings grown on Petri dishes. Extracts were incubated with μMACS anti-myc microbeads (Miltenyi Biotec, 130-091-123) for one hour at 4°C before recovering immune-complexes. For analysis of ING2-bound proteins from pull-down experiments, proteins were digested as described (Gomez-Zambrano et al., 2018), with minor modifications. After digestion, tryptic peptides were analysed by LC-MS/MS in an Orbitrap Fusion mass spectrometer (Thermo Fisher Scientific). An enhanced FT-resolution spectrum (resolution=70.000) followed by the HCD MS/MS spectra from the n-th most intense parent ions were analysed along the chromatographic run. Dynamic exclusion was set at 40 s.

For peptide identification, spectra were analysed with Proteome Discoverer v2.1.0.81 (Thermo Fisher Scientific) using SEQUEST-HT engine (Thermo Fisher Scientific). Database search was performed against the Uniprot database containing all sequences from *Arabidopsis thaliana* and crap contaminants (December 5th, 2016; 77084 sequences), using the following parameters: trypsin digestion with 2 maximum missed cleavage sites, precursor and fragment mass tolerances of 800 ppm and 1.2 Da respectively, carbamidomethyl cysteine as fixed modification and methionine oxidation as variable modifications. Scaffold program (version 5.3.3, Proteome Software Inc., Portland, OR) was used to validate MS/MS based peptide and protein identifications, with an additional filter for precursor mass tolerance of 15 ppm (Bonzon-Kulichenko et al., 2015). Protein probabilities were assigned by the Protein Prophet algorithm (Nesvizhskii et al., 2003). We performed two pull-down biological replicates and the corresponding control protein purification from WT, and the identified peptides are listed in Supplemental Table 1.

### Gene expression analysis

For RT-PCR and qPCR analysis, RNA was extracted from plants grown on the indicated conditions with the E.Z.N.A Plant RNA kit (Omega Bio-Tek). RNA was retro-transcribed using the Maxima first strand cDNA synthesis kit (Thermo Fisher Scientific). Quantitative PCR was performed using LightCycler 480 SYBR Green I (Roche). Primers used for qPCR analysis are listed on Supplementary Table 4. The *PP2AA3* (At1g13320) gene was used as a reference in all experiments.

For RNA sequencing experiments, 10 day-old seedlings were grown on petri dishes and plant tissue was harvested at ZT16. RNA was extracted from two independent experiments to prepare sequencing libraries for each genotype. RNA library preparation and sequencing was performed by Beijing Genome Institute (BGI) on a HiSeq2500 platform generating 4Gb clean 150PE reads per sample. Clean reads were mapped to Arabidopsis TAIR10 reference genome using HISAT2 v2.0.5 (Kim et al., 2015). Differential gene expression analysis was performed using the DESeq2 module from SeqMonk v1.35 software (Babraham Institute). Differentially expressed genes were defined by FDR ≤ 0.05 and log2 fold change ≥1 or ≤-1. Gene ontology (GO) enrichment analysis was performed on PANTHER (http://pantherdb.org/) using a Fisher’s exact test corrected by a false discovery rate FDR < 0.05 as cutoff for a significantly enriched GO term.

### Chromatin immunoprecipitation

For ChIP experiments we collected aerial tissue from soil-grown plants grown for 14 days at 22°C in LD conditions. Tissue was cross-linked with 1% formaldehyde for 10 min. Dual crosslinking was implemented for tissue from complemented myc-ING2 transgenic lines by incubating plant material for 30 min in the presence of 2 mM disuccinimidyl glutarate (DSG) followed by a 10 min incubation with 1% formaldehyde. Chromatin was isolated essentially as described (Komar et al., 2016). Histones were immunoprecipitated using the following antibodies: α-H4K5,8,12,16ac (Millipore, 06-598), α-H4 (Millipore, 05-858), α-H2A.Z (Agrisera, AS10718) while 4xmyc-ING2 was immunoprecipitated with, α-myc (Millipore, 05-724). For ChIP-qPCR experiments cross-links were reversed by a 10 min incubation at 95°C in the presence of 10% Chelex-100 resin (Biorad) followed by Proteinase-K treatment. For the ChIP-seq experiment we reversed cross-links by incubating chromatin overnight in 0.2 M NaCl at 65°C followed by Proteinase-K treatment. Libraries were prepared using QIAseq Ultralow input library kit (QIAgen) and sequencing was performed by Novogene (https://www.novogene.com) in a Novaseq X Plus instrument generating 6 Gb of 150PE reads. Clean reads were mapped to Arabidopsis TAIR10 reference genome using bowtie2 v2.3.5.1. Metaplots were generated using deepTools v3.5.4. Enriched peaks were called using MACS2 v2.2.6 and differential peak analysis was performed using the bdgdiff module from this program.

## Accession numbers

The complete sequencing data from this publication were submitted to the Gene Expression Omnibus database (www.ncbi.nlm.nih.gov/geo/) under accession numbers xxx (RNA-seq) and xxxx (ChIP-seq).

## Acknowledgements

This work is supported by competitive grants PID2022-137131NB-I00 to J.A.J. and M.P. and PID2021-122348NB-I00 funded by MCIN/AEI/10.13039/501100011033/FEDER, UE; and by “the Comunidad de Madrid grant S2022/BMD-7333-CM and la Caixa Foundation grant LCF/PR/HR22/52420019. A.M. was granted with a FPU fellowship from the Spanish Ministry of Education and Y.T. is recipient of the Chinese Scholarship Council (CSC) predoctoral fellowship 202109210045. The CBGP is a Severo Ochoa Center of Excellence (grant CEX2020-000999-S funded by MICIU/AEI/10.13039/501100011033). The CNIC is supported by the Instituto de Salud Carlos III (ISCIII), the Ministerio de Ciencia, Innovación Y Universidades (MICIU) and the Pro CNIC Foundation), and is a Severo Ochoa Center of Excellence (grant CEX2020-001041-S funded by MICIU/AEI/10.13039/501100011033).

## Author contributions

J.A.J and M.P conceived and supervised the research. J.B-G carried out the ChIP-seq and ChIP-PCR experiments, and together with A.M. and Y.T performed the physiological, molecular and genetic analyses shown, as well as the RNA-seq data. R.P and P.C contributed with the preparation of the ING2-pull-down samples for AP/MS while J.A.L and J. V. carried out the proteomic analysis of them, and analyzed the resulting data. J.B-G wrote the manuscript with direct inputs from J.A.J and M.P.

## Competing interests

The authors declare no competing interests.

---

**Supplemental Figure 1:**
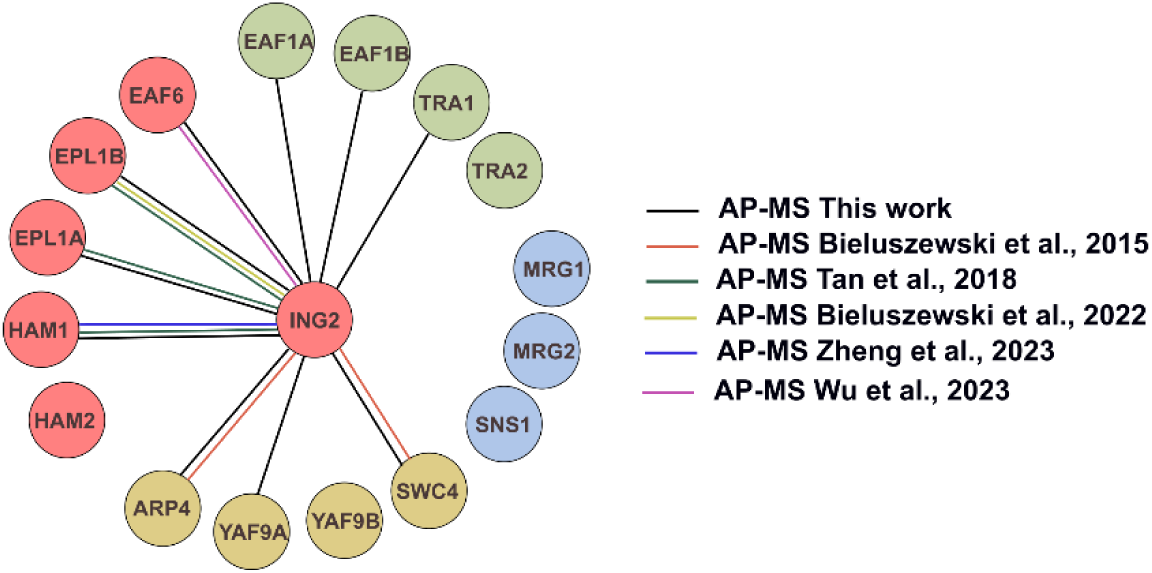
Interaction of ING2 with NuA4-C subunits. Schematic diagram showing current AP-MS evidence supporting ING2 interaction with NuA4-C subunits. Circle colors indicate NuA4-C modules: piccolo (red), core (green), TINTIN module (blue), and YEATS (ochre). Line colors indicate AP-MS experiments and the corresponding references.

**Supplemental Figure 2:**
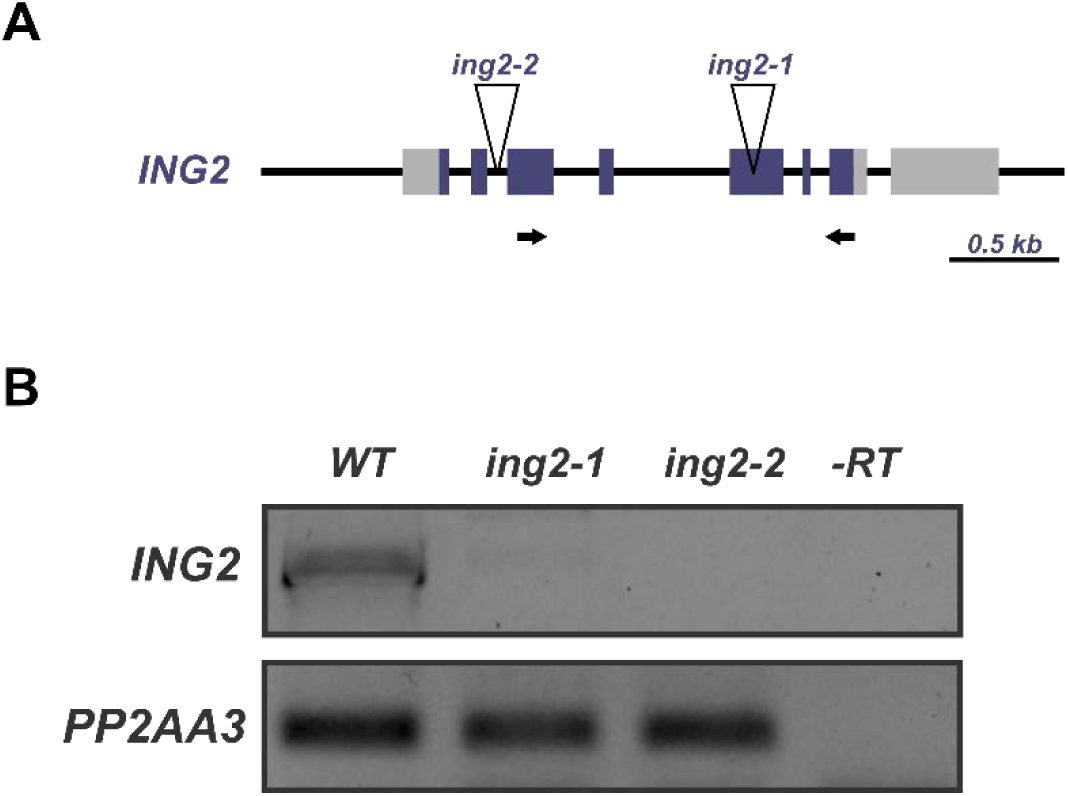
Validation of *ING2* T-DNA insertion lines used in this study. (A) Schematic representation of the *ING2* gene showing the location of T-DNA insertions in *ing2-1* and *ing2-2* mutant lines. Primers used to check *ING2* expression in T-DNA lines are indicated by arrows. (B) RT-PCR gene expression analysis in plants from the indicated genotypes. As a negative control (-RT) we used a cDNA synthesis product lacking retro transcriptase enzyme.

**Supplemental Figure 3:**
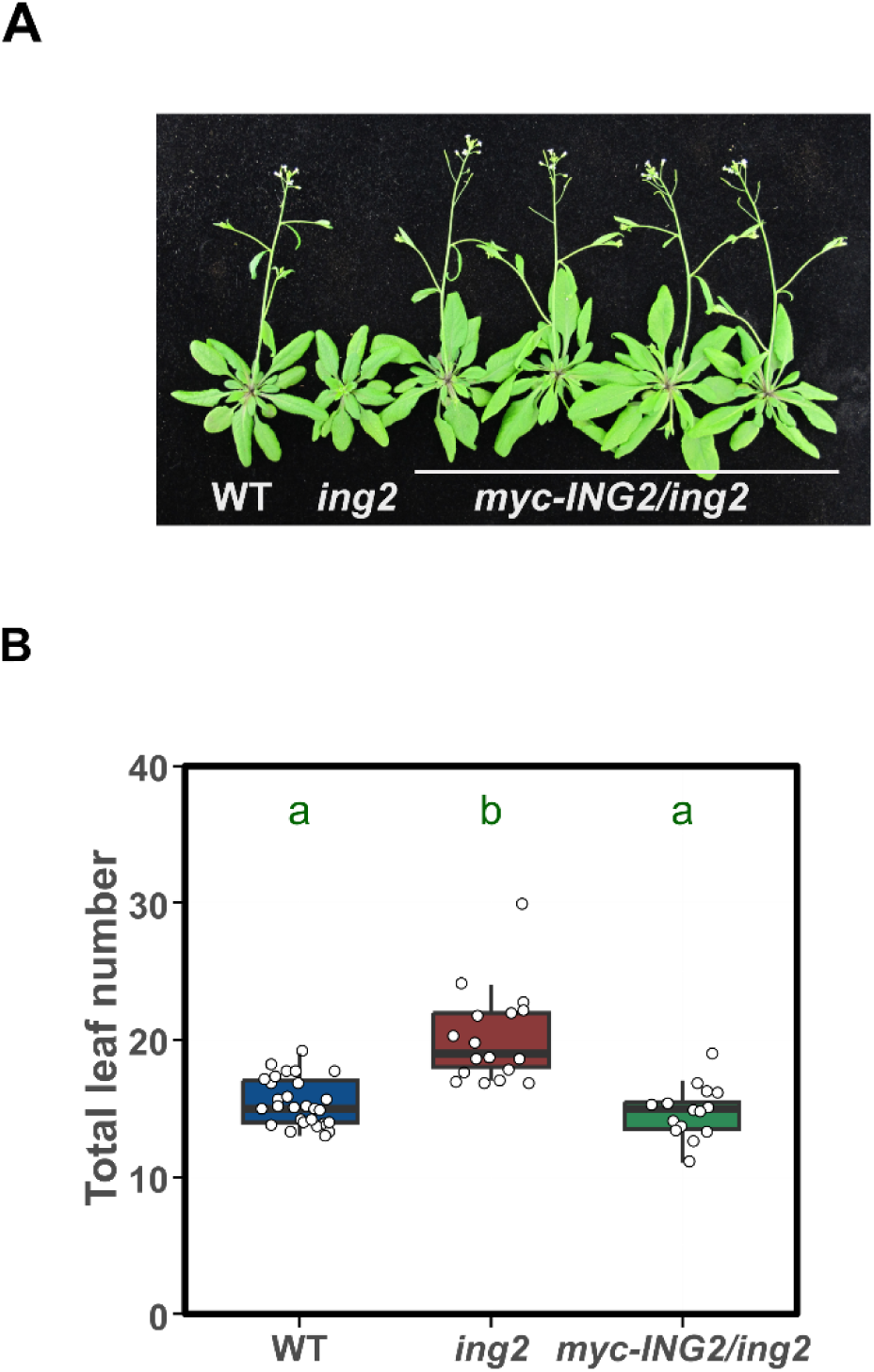
A myc-tagged ING2 construct complements the late flowering displayed by the *ing2-2* mutant. (A) Representative plants of the indicated genotypes grown under LD conditions. (B) Determination of flowering time in the indicated genotypes. Statistical significance in a one-way ANOVA test is indicated by different letters.

**Supplemental Figure 4:**
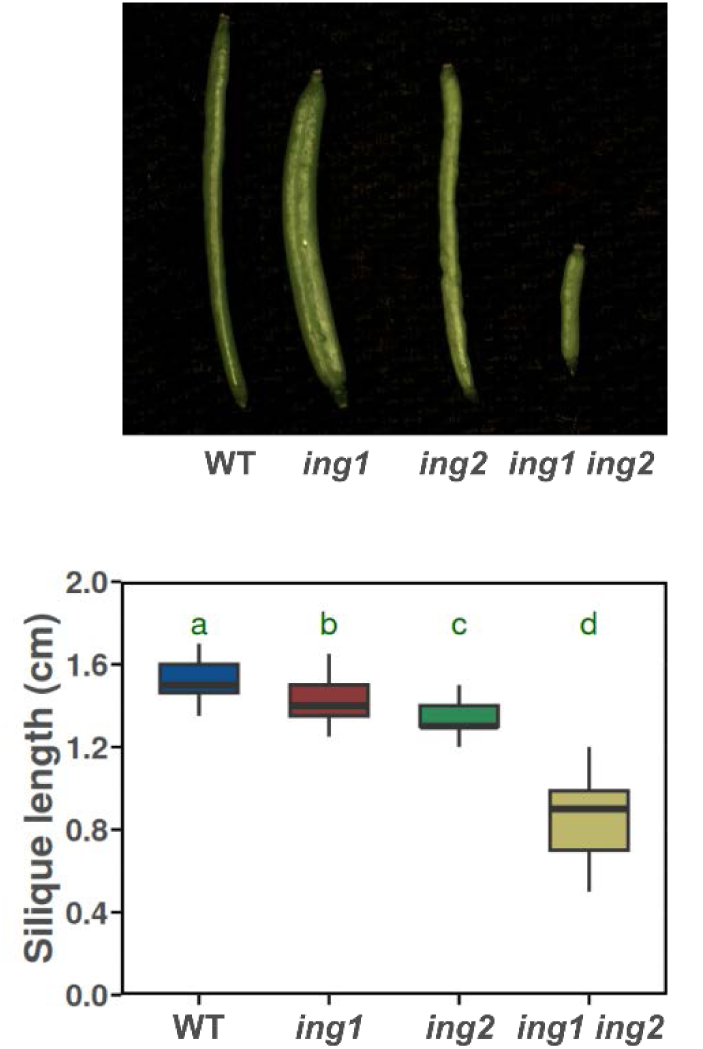
*ing1 ing2* double mutants cause an enhancement of fruit developmental defects. Fruit length in LD grown plants of the indicated genotypes. Statistical significance in a one-way ANOVA test is indicated by different letters.

**Supplemental Figure 5:**
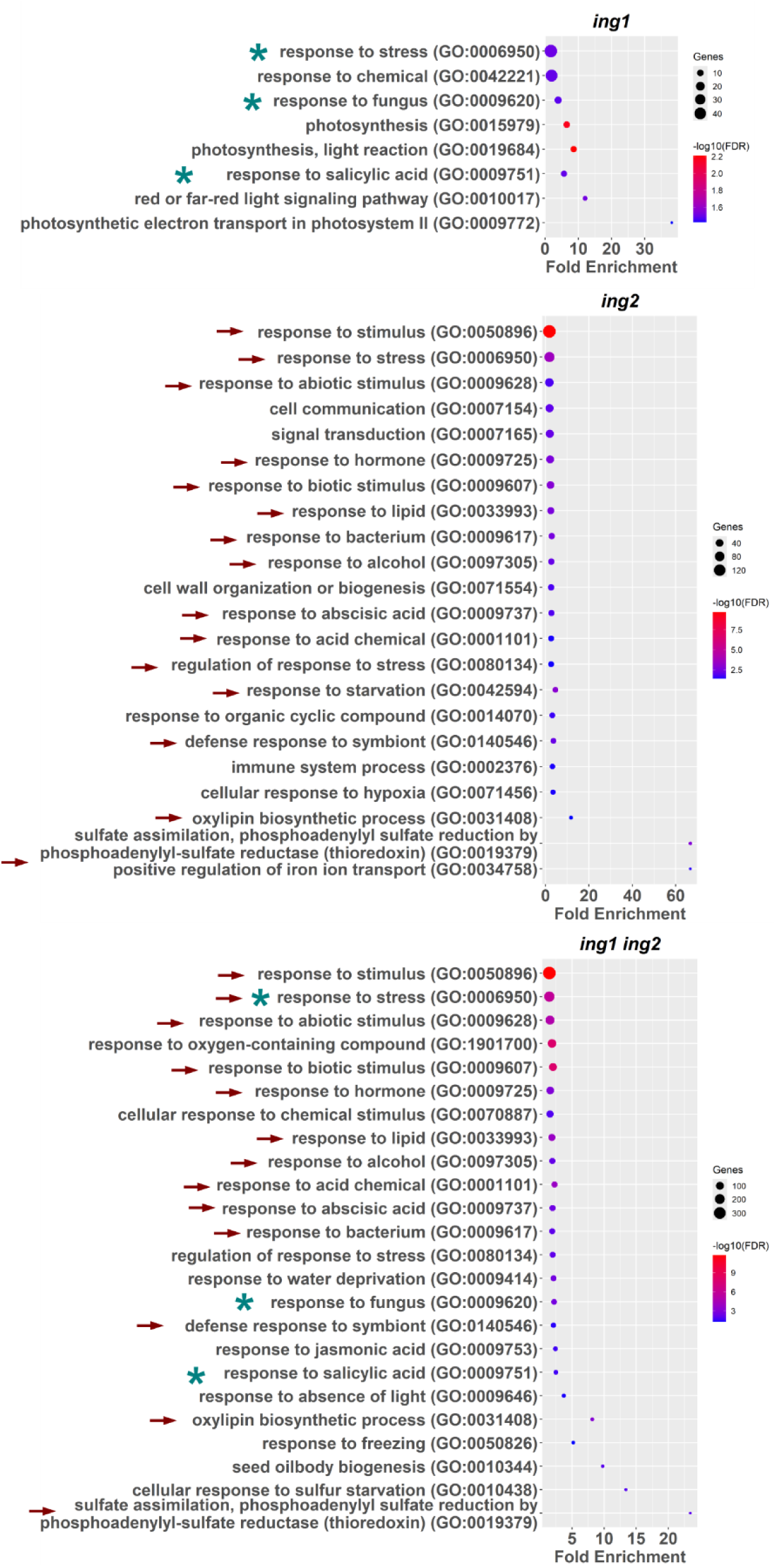
Gene ontology enrichment analysis on differentially expressed genes in *ing* mutants. Selected enriched GO terms (biological processes category) among identified DEGs in *ing* mutants. Shared GO terms between *ing1 ing2* and *ing1* (blue asterisks) or *ing2* mutants (red arrows) are indicated.

**Supplemental Figure 6.**
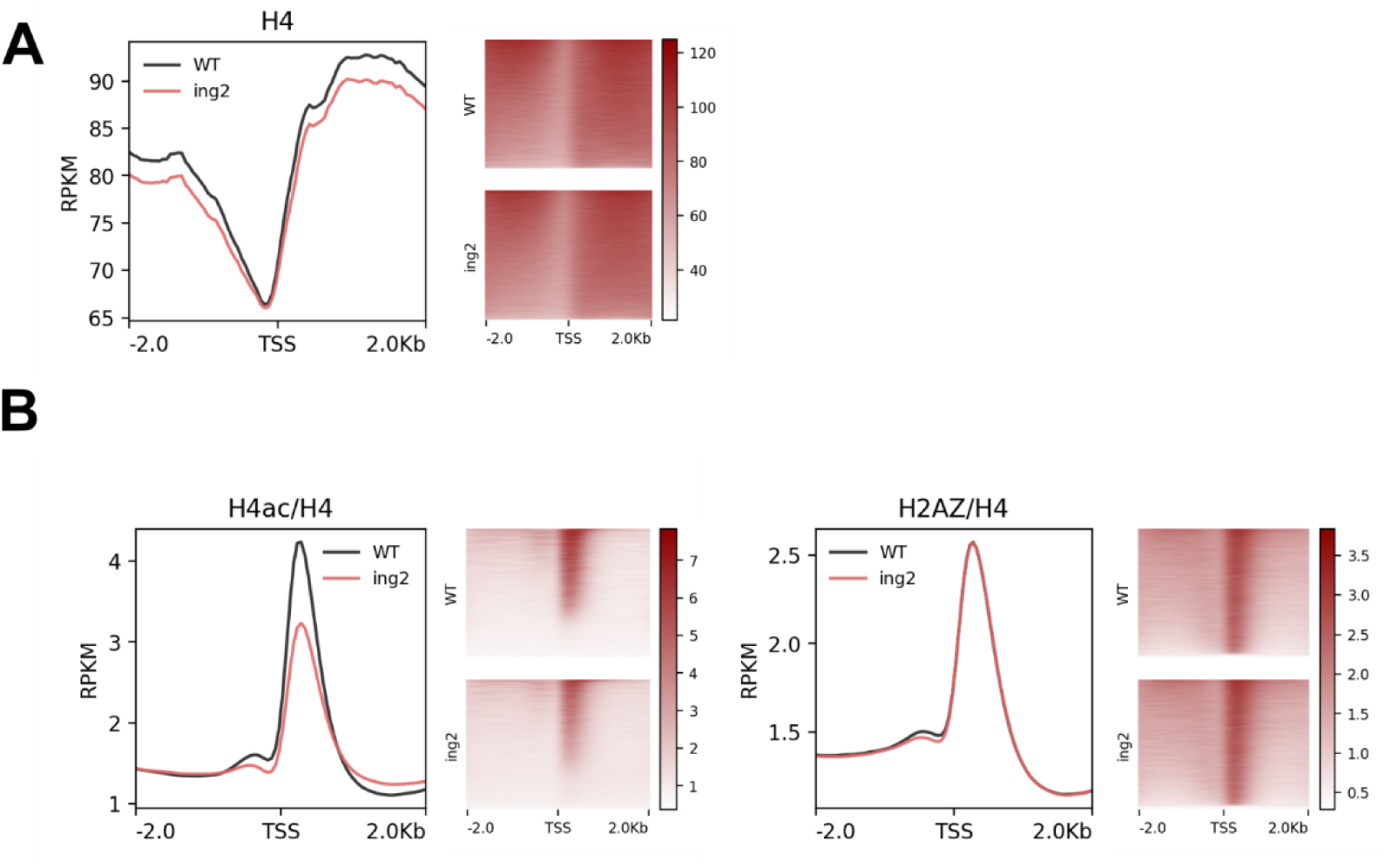
Differences in H4 occupancy do not explain the lower levels of H4ac found in the *ing2* mutant. (A) TSS-centered profile and read density map displaying H4 occupancy levels in WT and *ing2* mutant. (B) Metaplots and read density maps showing H4-normalized levels of H4ac and H2A.Z in WT and *ing2* mutant.

## Notes

### Competing Interest Statement

The authors have declared no competing interest.

